# Assembly and Activation of Dynein-Dynactin by the Cargo Adaptor Protein Hook3

**DOI:** 10.1101/047605

**Authors:** Courtney M. Schroeder, Ronald D. Vale

## Abstract

Metazoan cytoplasmic dynein moves processively along microtubules with the aid of dynactin and an adaptor protein that joins dynein and dynactin into a stable ternary complex. Here, we have examined how Hook3, a cargo adaptor involved in Golgi and endosome transport, forms a motile dynein-dynactin complex. We show that the conserved Hook domain interacts directly with the dynein light intermediate chain 1 (LIC1). By solving the crystal structure of the Hook domain and using structure-based mutagenesis, we identify two conserved surface residues that are each critical for LIC1 binding. Hook proteins with mutations in these residues fail to form a stable dynein-dynactin complex, revealing a crucial role for LIC1 in this interaction. We also identify a region of Hook3 specifically required for an allosteric activation of processive motility. Our work reveals the structural details of Hook3’s interaction with dynein and offers insight into how cargo adaptors form processive dynein-dynactin motor complexes.

## Introduction

Eukaryotic cells use molecular motors to transport and spatially organize organelles, proteins, and mRNAs within the cytoplasm. Cytoplasmic dynein is a molecular motor that carries cargo towards microtubule minus ends (reviewed in Allan, 2011). Dynein is a large homodimer composed of two ~500 kDa heavy chains that contain the ATPase motor domain (reviewed in Bhabha et al., 2016, Schmidt et al., 2015). The N-terminal portion of the heavy chain binds additional subunits known as the dynein “tail” subunits, which include the light intermediate chain (LIC), the intermediate chain (IC) and the light chains (LCs- LC8, Tctex1 and LC7/ roadblock, Pfister and Lo, 2012; Pfister et al., 2006)). This tail complex is responsible for linking dynein to cargo (reviewed in Pfister and Lo, 2012). Mammalian dynein is not constitutively active, but rather its motility is regulated by cargo interaction (McKenney et al., 2014; Schlager et al., 2014).

The mammalian light intermediate chains, encoded by two closely related gene products LIC1 and LIC2 (Hughes et al., 1995; Tynan et al., 2000), are involved in several different types of cargo interactions and dynein-based movements, including endosomal and lysosomal transport, ER export, Golgi transport, and axonal vesicle trafficking (Palmer et al., 2009; Horgan et al., 2010; Kong et al., 2013; Tan et al., 2011; Brown et al., 2014; Koushika et al., 2004). The domain structure of the LIC allows it to interact with cargo adaptors while integrated into the dynein holoenzyme. The LIC’s highly conserved N-terminal G protein-like domain binds directly to the dynein heavy chain and the less conserved C-terminal domain binds adaptor proteins (Schroeder et al., 2014, Fig. 1 A). These cargo adaptors are themselves multi-functional proteins that can bind to a protein (e.g. a Rab GTPase) on a membranous cargo (reviewed in Carter et al., 2016; Cianfrocco et al., 2015; Fu and Holzbaur, 2014).

**Figure 1.**
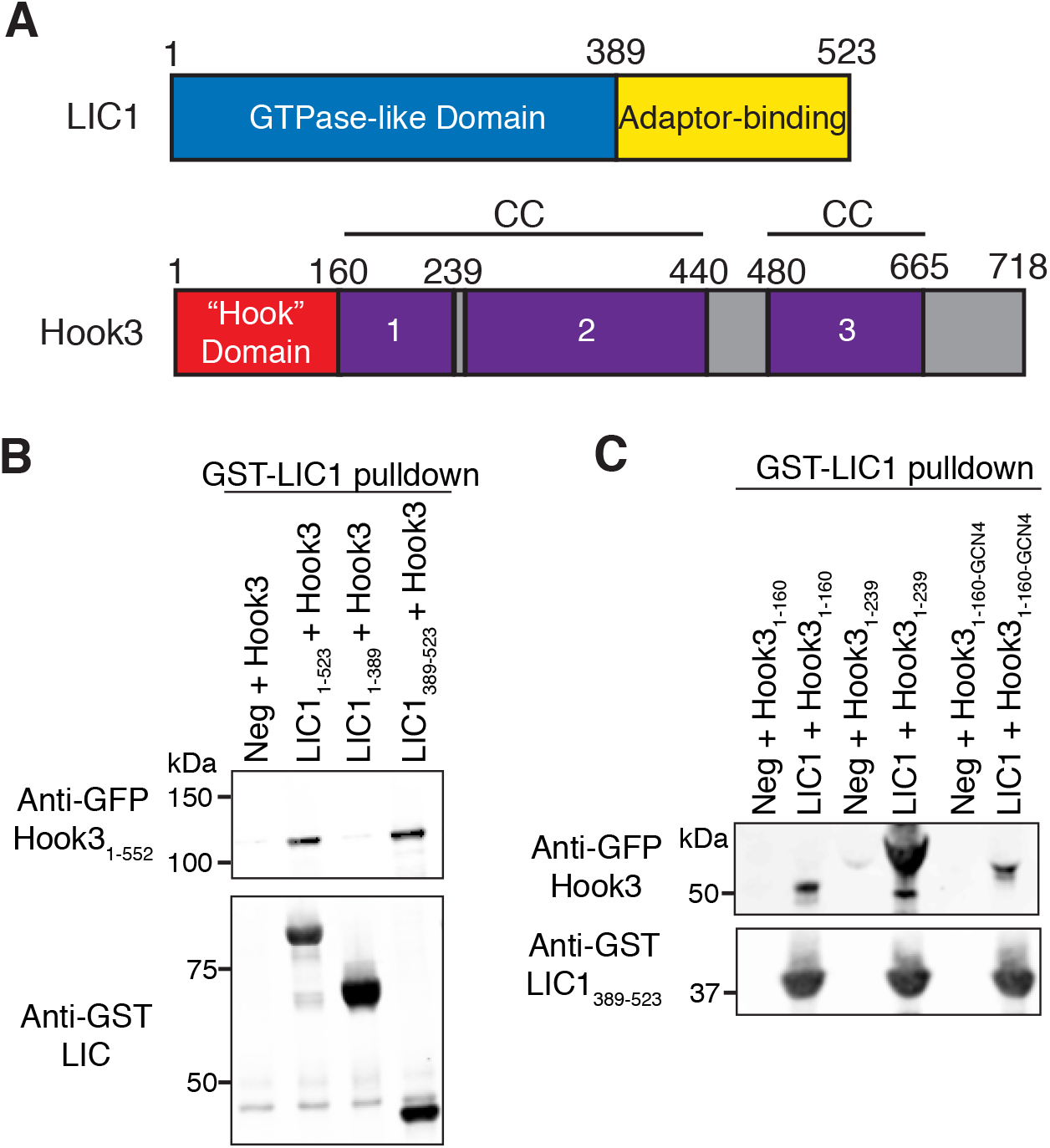
**Dynein LIC binds the “Hook domain” A)** The domain architectures of human LIC1 and Hook3. See text for details. **B)** GST-tagged human full-length or truncated LIC1 bound to glutathione resin were incubated with sfGFP-tagged Hook3_1-552_, centrifuged, and probed for Hook3 with an anti-GFP antibody. LIC1 in the pelleted beads was detected using an anti-GST antibody. Negative (Neg) control lacks LIC1 on the beads. **C)** Two sfGFP-tagged Hook3 constructs were tested for LIC1 binding using the assay described in (B). Also tested was a Hook domain artificially dimerized using a GCN4 sequence (Hook3_1-160-GCN4_). Negative control lacks LIC1 on the beads. Results are representative of two experiments performed on separate days.

In addition to binding dynein LIC and cargo, adaptor proteins also promote an interaction between dynein and dynactin, a 12 subunit protein complex (Schroer, 2004). For mammalian dynein, the formation of this tripartite complex is important for long distance movement (processivity) along microtubules (McKenney et al., 2014; Schlager et al., 2014). This mechanism has been best studied for Bicaudal D2 (BicD2), an adaptor that links dynein-dynactin to Rab6 on Golgi-derived vesicles (reviewed in (Dienstbier and Li, 2009). The N-terminus of BicD2 consists of a 270-residue coiled coil that sits in a groove of the Arp1 filament of dynactin and also interacts with the N-terminal region of the dynein heavy chain; this dynein heavy chain-BicD2-Arp1 interaction was proposed to stabilize the tripartite complex (Urnavicius et al., 2015). How this interaction promotes motility is less clear. One possibility is that cargo adaptors activate an autoinihibited state of dynactin, enabling it to bind to microtubules and help to initiate motility. Urnvicius et al. (2015) also suggested that cargo adaptors and dynactin reposition the motor domains of the dynein dimers (see also Torisawa et al., 2014). These models, however, have not proposed a role for a LIC-adaptor protein interaction. Furthermore, it is unclear whether the assembly of the tripartite motor complex and activation of motility are separable functions.

One cargo adaptor that has been shown to assemble and activate dynein-dynactin is Hook3, although its mechanism has been less studied compared to BicD2. The Hook proteins, first identified for their role in endosomal trafficking in *Drosophila* (Krämer and Phistry et al., 1999), are a widely expressed class of dynein-associated cargo adaptor proteins. *Drosophila* and fungi have a single Hook gene, while mammals have three Hook genes. The most conserved region of the Hook genes is found at the N-terminal domain (aa 1~164). Without this “Hook” domain, Hook can no longer interact with dynein and dynactin (Zhang et al., 2014; Bielska et al., 2014). Following the N-terminal domain are three coiled coil domains that are important for dimerization (Krämer and Phistry et al., 1999; Walenta et al., 2001), and a divergent C-terminal domain (~aa 552-718) binds a variety of proteins that are specific for each Hook isoform (Maldonado-Báez et al., 2013; Szebenyi et al., 2006; Moynihan et al., 2009; Sano et al., 2007; Walenta et al., 2001). All mammalian Hook isoforms promote endosomal trafficking by forming a complex with Fused Toes (FTS) and Hook Interacting Protein (FHIP) (Xu et al., 2008).

Here, we sought to understand the mechanism by which Hook3 interacts with dynein and dynactin and activates processive motility. We report the crystal structure of the Hook domain and show that this domain binds directly to the C-terminal region of LIC1. Structure-based mutagenesis studies revealed two conserved surface residues that are essential for this interaction. Abrogation of the LIC interaction renders Hook3 unable to join dynein and dynactin into a stable complex. Interestingly, while the N-terminal 239 residues of Hook3 are sufficient for forming a stable complex with dynein-dynactin, this tripartite complex is immotile; activation of motility requires a more distal coiled coil region of Hook3. This result reveals that complex assembly and the activation of motility are separable activities. Our data suggest a model for how Hook3 joins dynein and dynactin into a motile complex.

## Results

### The Hook domain of Hook3 binds to the dynein LIC

Hook3 is comprised of the N-terminal, highly conserved “Hook” domain (Walenta et al., 2001), followed by three coiled coils and a C-terminal cargo-binding region (Fig. 1 A). A yeast two-hybrid assay revealed an interaction between aa 1-236 of *C. elegans* Hook and the LIC (Malone et al., 2003). We sought to confirm a direct interaction between Hook3 and LIC1 using purified proteins, as we demonstrated previously for the adaptor proteins RILP, BicD2, and FIP3 (Schroeder et al., 2014). Previous work showed that Hook3_1-552_ is sufficient to produce a highly processive dynein-dynactin-Hook3 complex (McKenney et al., 2014), and thus we used this slightly truncated Hook3 protein to test for interactions with LIC1. GFP-tagged Hook3_1-552_ was incubated with beads coated with GST-tagged versions of either full-length LIC1, the LIC N-terminal G-domain (LIC1_1-389_) or the C-terminal domain (LIC1_389-523_); the beads and any interacting proteins were centrifuged and the protein composition of the pulldown was analyzed by immunoblot. The results revealed that Hook3_1-552_ cosedimented with full-length LIC1 and the LIC1 C-terminus alone, but not with the N-terminal LIC1 G domain (Fig. 1 B, Fig. S1 A). Thus, similar to the other cargo adaptors RILP, BicD2 and FIP (Schroeder et al., 2014), Hook3 also directly binds to LIC1_389-523_.

We next truncated Hook3_1-552_ to identify a smaller fragment that might bind LIC1_389-523_. The shorter truncation Hook3_1-239_ bound to LIC1_389-523_ in the pulldown assay (Fig. 1 C, Fig. S1 B) and the two proteins co-eluted as a stable complex by gel filtration chromatography (Fig. S1 C). The Hook domain alone (Hook3_1-160_) also bound LIC1_389-523_, albeit more weakly than Hook3_1-239_ (Fig. 1 C, Fig. S1 B). Hook3_1-160_ lacks the predicted coiled coil found in Hook3_1-239_ and thus the stronger interaction of Hook3_1-239_ might be due to the fact that it is a dimer. We therefore tested an artificially dimerized coiled coil version of Hook3_1-160_ (Hook3_1-160-GCN4_) but found that its binding affinity to LIC1 was not increased relative to the monomeric version (Fig. 1 C, Fig. S1 B). Overall, these data indicate that the N-terminal Hook domain can bind specifically to the C-terminal region of LIC1 and that the region between aa 160-239 strengthens this interaction.

### The Hook domain contains a calponin homology fold with an extended α-helix

We attempted to co-crystallize LIC1_389-523_ with either Hook3_1-239_ or Hook3_1-160_, but crystals were only obtained for Hook3_1-160_ alone. A 1.7 Å dataset was obtained from one of the crystals, and a poly-alanine model based upon an NMR solution structure of mouse Hook1_1-160_ (PDB 1WIX) was used for molecular replacement. After multiple rounds of refinement, the final structure has an R_work_ of 18.4 and R_free_ of 21.9 (Table I). Two copies of the protein are present in the asymmetric unit and interact through an anti-parallel arrangement of their C-terminal α-helices (helix H described below). This interaction may not be physiological since Hook3_1-160_ is monomeric, as determined by static light scattering (data not shown).

**Table I.**
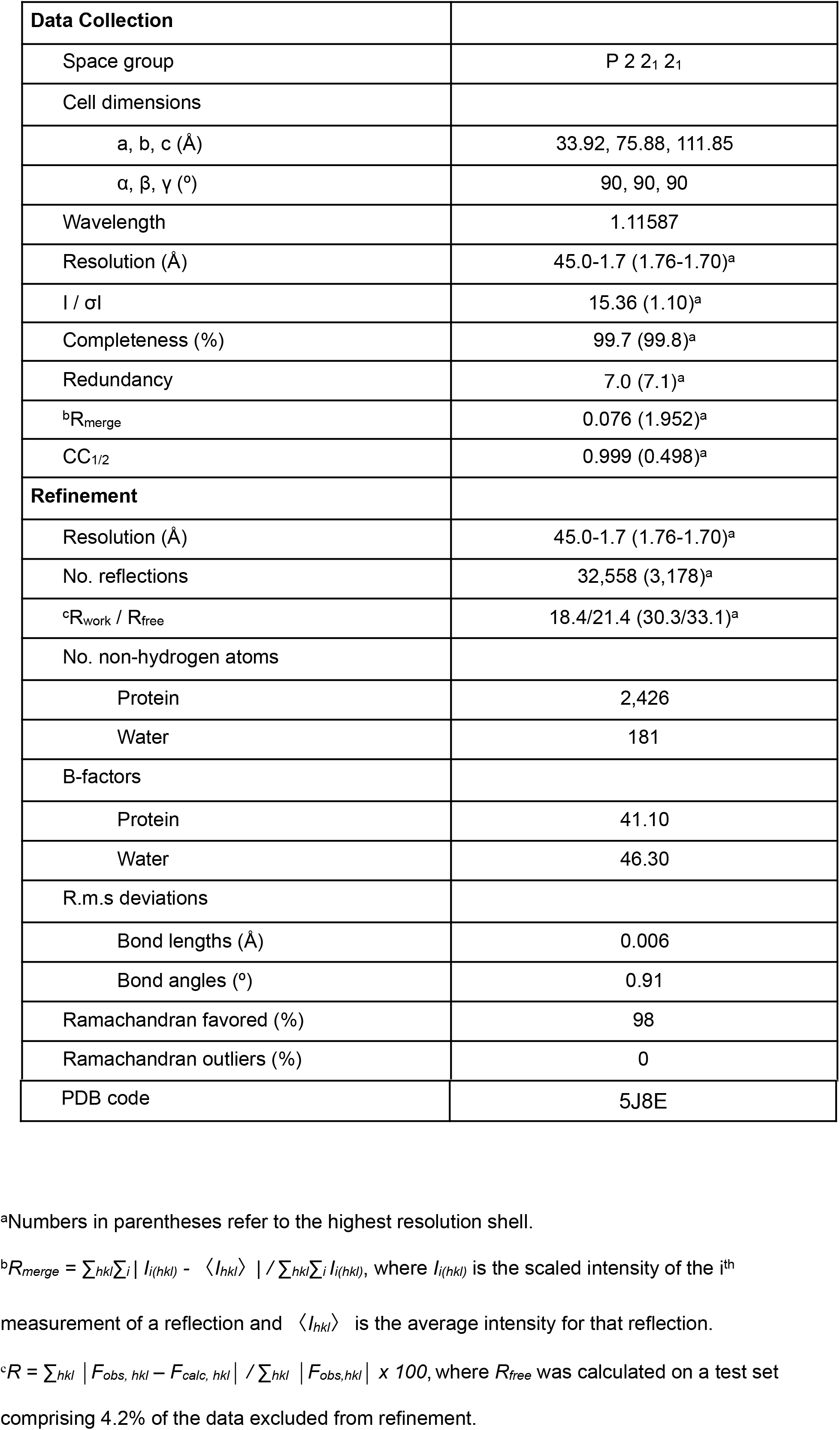
**Crystallographic Data and Refinement Statistics**

The Protein Homology/analogY Recognition Engine (PHYRE), which predicts a protein’s tertiary structure based on homology, previously predicted that the Hook domain is comprised of a calponin homology (CH) fold (Zhang et al., 2014). Our structure indeed exhibits a canonical 7-helix CH fold (Fig. 2 A). However, the crystal structure reveals an additional 8^th^ α-helix (helix H, aa 132-158; Fig. 2 A) extending at ~90° from helix G, which was not expected from prior secondary structure prediction (Drozdetskiy et al., 2015). This same α-helix also appears in the NMR structure of the Hook domain of mouse Hook1 (PDB 1WIX), but it is bent in the middle and folded back on itself (Fig. S2 A). Thus, it appears that helix H is able to adopt different conformations; the extended conformation that we have observed may be stabilized by proteinprotein interactions in the asymmetric unit.

**Figure 2.**
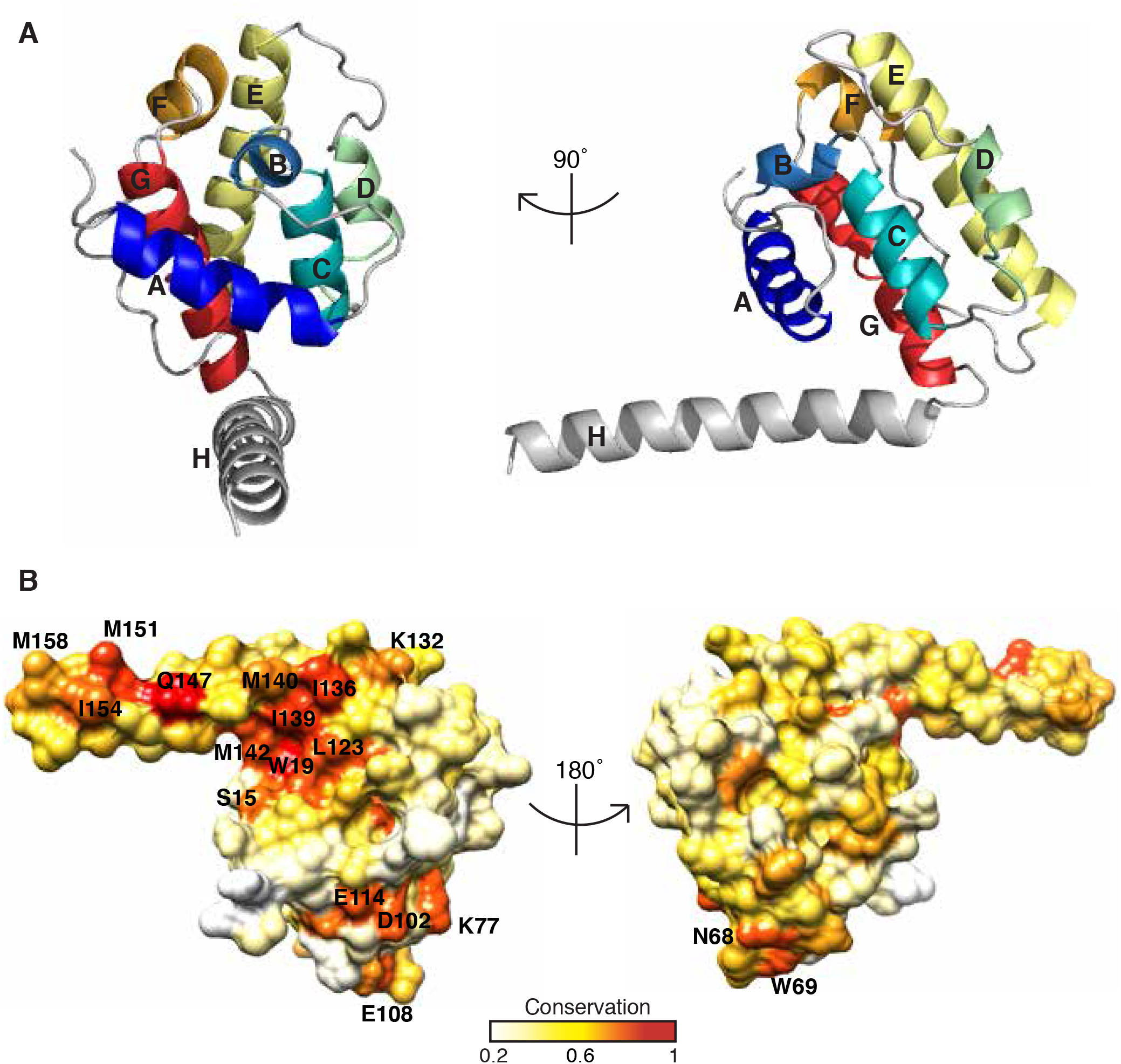
**The structure of the Hook domain exhibits an extended α-helix and restricted conservation. A)** The 1.7 Å structure of the Hook domain (aa 9-158) from human Hook3 with the helices labeled A-H. **B)** The conservation of residues on the surface of the structure in (A) is shown with red representing the most conserved and white depicting the least conserved. Highly conserved residues are labeled.

We next mapped the conserved surface residues in the Hook domain onto our crystal structure using an alignment of 19 Hook domain sequences ranging from fungal to mammalian species (Fig. S3). Strikingly, one side of the structure is much more highly conserved than the other (Fig. 2 B). This contrast is even more evident in the map of conserved residues between the three human Hook genes (Fig. S2 B). Several highly conserved residues lie within the extended helix H, including the universally conserved Q147 and nearby conserved hydrophobic residues. Two other prominent patches of conservation lie on this same face of the CH domain— one cluster consists mainly of hydrophobic residues (S15, W19, L121, L123), and the other consists of charged residues (K77, D102, E108 and E114).

### Two Highly Conserved Residues Mediate the Hook3-Dynein Interaction and are Critical for Dynein-Dynactin Motility

The surface conserved residues could be part of a binding interface with the dynein light intermediate chain. To test which region of the Hook domain might be involved in binding LIC1, we made four proteins with different clusters of alanine mutants: 1) Q147A/M151A/I154A, 2) I136A/I139A/M142A, 3) N68A/W69A/K77A and 4) D102A/E108A (Fig. 3 A). These mutations were made in the construct Hook3_1-239_ due to its higher binding affinity to LIC1 than Hook3_1-160_. The triple and double mutations produced monodisperse protein, as assessed by gel filtration (Fig. S4 A). We tested each mutant Hook3 protein for binding to GST-LIC1_389-523_ using the bead pulldown assay (Fig. 3 B, Fig. S4 B). The triple mutants Q147A/M151A/I154A and I136A/I139A/ M142A exhibited little or no detectable binding. In contrast, the triple mutant N68A/W69A/K77A and the double mutant D102A/E108A showed little difference in binding (Fig. 3 B, Fig. S4 B). Since patches Q147A/M151A/I154A and I136A/I139A/M142A both lie within helix H, these results suggest the highly conserved helix H contains the main LIC1 binding interface.

**Figure 3.**
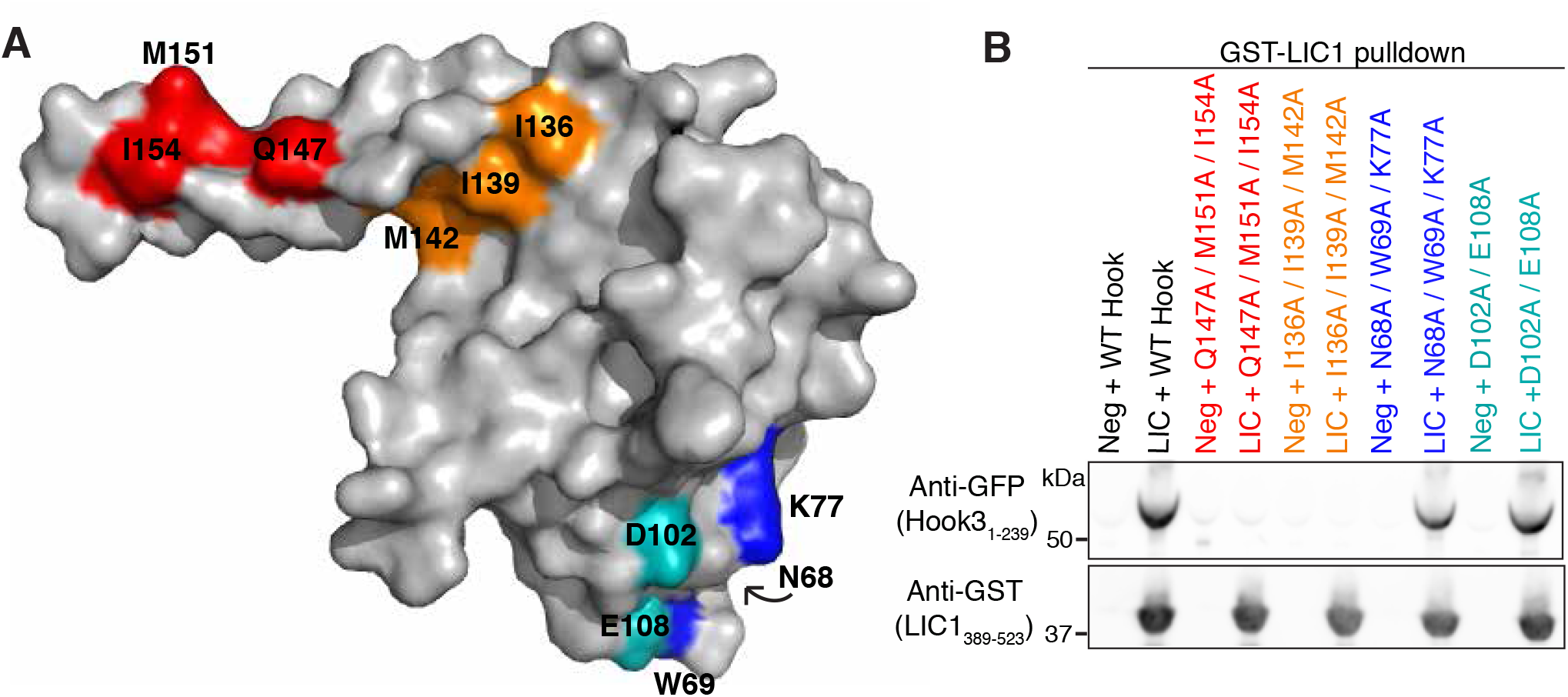
**Helix H in the Hook domain contains a LIC-binding interface A)** Patches of conserved residues in the Hook domain were mutated in separate constructs. Each patch of residues is denoted by a different color. **B)** GST-LIC1_389-523_, bound to glutathione resin, was incubated with sfGFP-Hook3_1-239_ mutants; the beads were centrifuged and then Hook3 binding was assessed by immunoblot analysis using an anti-GFP antibody. The presence of the bait GST-LIC1_389-523_ was verified using an anti-GST antibody. Negative control lacks LIC1 on the beads. Blots are representative of two independent experiments performed on separate days.

We next investigated the more solvent-exposed Q147A/M151A/I154A patch in more depth with point mutants. Strikingly, the single I154A and Q147A mutations each led to a dramatic reduction in the Hook3-LIC1 interaction (Fig. 4A, Fig. S4 C). In contrast, the Hook3 mutant M151A could still bind LIC1 as well as wild-type. (Fig. 4 A, Fig. S4 C). We next tested whether these single point mutants affected the binding of Hook3_1-552_ to intact dynein and dynactin in a porcine brain lysate (McKenney et al., 2014). After adding wild-type or Hook3 mutants to the lysate followed by the Hook3 pulldown, we found that the single point mutants Q147A and I154A bound very little or no dynein and no detectable dynactin, whereas the M151A mutant bound dynein-dynactin in a similar manner to wild-type (Fig. 4 B, Fig. S4 D). These results indicate that the highly conserved residues Q147 and I154 in helix H of Hook3 both play critical roles in binding LIC1 and in forming a stable dynein-dynactin complex.

**Figure 4.**
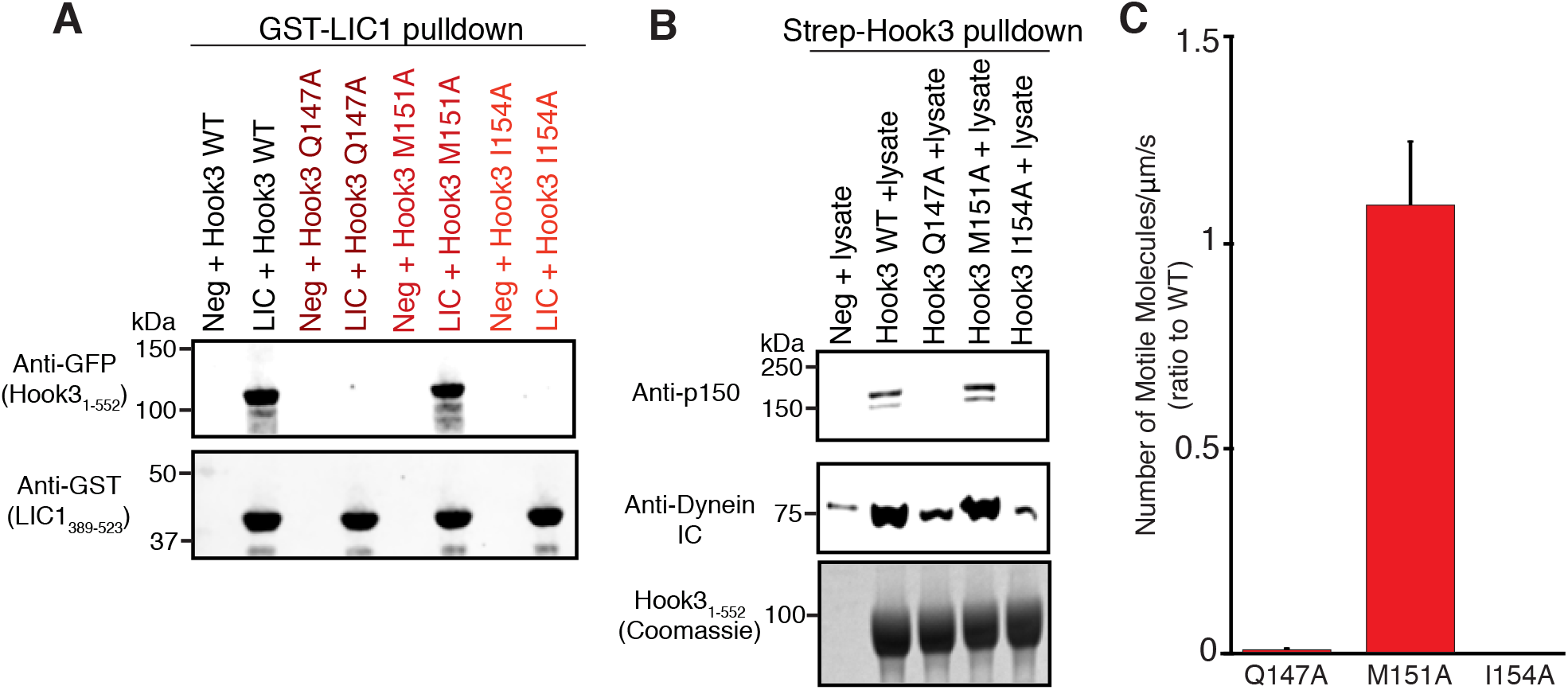
**Two conserved Hook3 residues are critical for the assembly and motility of dynein-dynactin A)** Single point mutations Q147A, M151A, and I154A in sfGFP-tagged Hook3_1-552_ were compared to wild-type (WT) and tested for binding to human GST-LIC1_389-523_ as in Fig. 3 B (representative of triplicate experiments). Negative control lacks LIC1 on the beads. **B)** StrepII-Hook3 constructs, bound to Strep-Tactin resin, were incubated with porcine brain lysate, and then the beads were centrifuged and the resin analyzed by immunoblot for the dynein intermediate chain (IC) and the p150 subunit of dynactin. Negative control lacks Hook3 on the beads. The amount of each Hook3 construct was assessed by Coomassie stain. Representative of two independent experiments. **C)** The wild-type (WT) and single point mutants were incubated with affinity-purified human dynein-dynactin and 1 mM ATP. SfGFP-tagged Hook3_1-552_ was visualized by TIRF microscopy and classified as “processive” if it moved unidirectionally for >1 µm along microtubules. All constructs were normalized by dividing the total number of processive motors by the total length of microtubules in the field of view and the time of the movie (movements/µm/s). The ratios of the mutants to wild-type were calculated from side-by-side experiments performed on the same day. Shown are the mean ± SD of the ratios from two independent experiments performed on different days.

We next tested the ability of Hook3 mutant proteins to stimulate dynein-dynactin motility (McKenney et al., 2014). Dynein and dynactin, purified from a human RPE-1 cell line, were pre-incubated with GFP-tagged Hook3 constructs. The mixture was then added in the presence of ATP to glass-immobilized microtubules and interactions of GFP-Hook3 with microtubules were examined by total internal reflection fluorescence (TIRF) microscopy. Processive movement of dynein-dynactin and wild-type GFP-Hook3 was observed as previously described. The point mutant M151A produced a similar number of motile dynein-dynactin-Hook3 molecules compared to wild-type GFP-Hook3 (Fig. 4 C), and the velocities of the molecules were in a similar range as wild-type Hook3 (Fig. S5 E). In contrast, Q147A and I154A GFP-Hook3 constructs did not elicit processive runs (Fig. 4 C, Fig. S4 E), presumably because they did not bind to and form a complex with dynein and dynactin. Thus, Q147 and I154 are each essential for Hook3’s interaction with LIC1 and for the formation of a processive dynein-dynactin complex.

### Hook3 truncations that assemble dynein-dynactin do not elicit processive motility

We next sought to define the roles that the Hook domain and the extended coiled coil domains of Hook3 play in assembling dynein and dynactin into a complex. Prior work on the 270-residue coiled coil domain of BicD2 showed that it sits in the groove of the dynactin Arp1 filament and creates a binding interface with the dynein heavy chain (Urnavicius et al., 2015). We made two constructs that consisted primarily of the Hook domain (aa 1-160 and 1-239), truncations that excluded the Hook domain (aa 160-552, and 239-552), and a truncation that excluded just the CH domain but contained helix H of the Hook domain (aa 130-552) (Fig. 1 A). These constructs, bound to Strep-Tactin resin, were incubated with porcine brain lysate and then assessed for their ability to pulldown the endogenous dynein-dynactin complex by immunoblotting for the dynein IC and the dynactin subunit p150. The construct Hook3_1-239_ pulled down both dynein and dynactin, albeit to a lesser extent than the longer Hook3_1-552_ (Fig. 5 A, Fig. S5 A). Hook3_1-160_ pulled down a small amount of dynein, but the dynactin signal appeared equivalent to that of the negative control (Fig. 5 A, Fig. S5 A). The relative amounts of dynein pulled down by Hook3_1-160_ and Hook3_1-239_ are analogous to the relative binding affinities for purified LIC1 (Fig. 1 C). The constructs lacking the Hook domain did not pulldown dynein or dynactin, similar to what was found *in vivo* for HookA in *Aspergillus nidulans* (Zhang et al., 2014). Overall, these data demonstrate the importance of the Hook domain for the formation of this tripartite motor complex.

**Figure 5.**
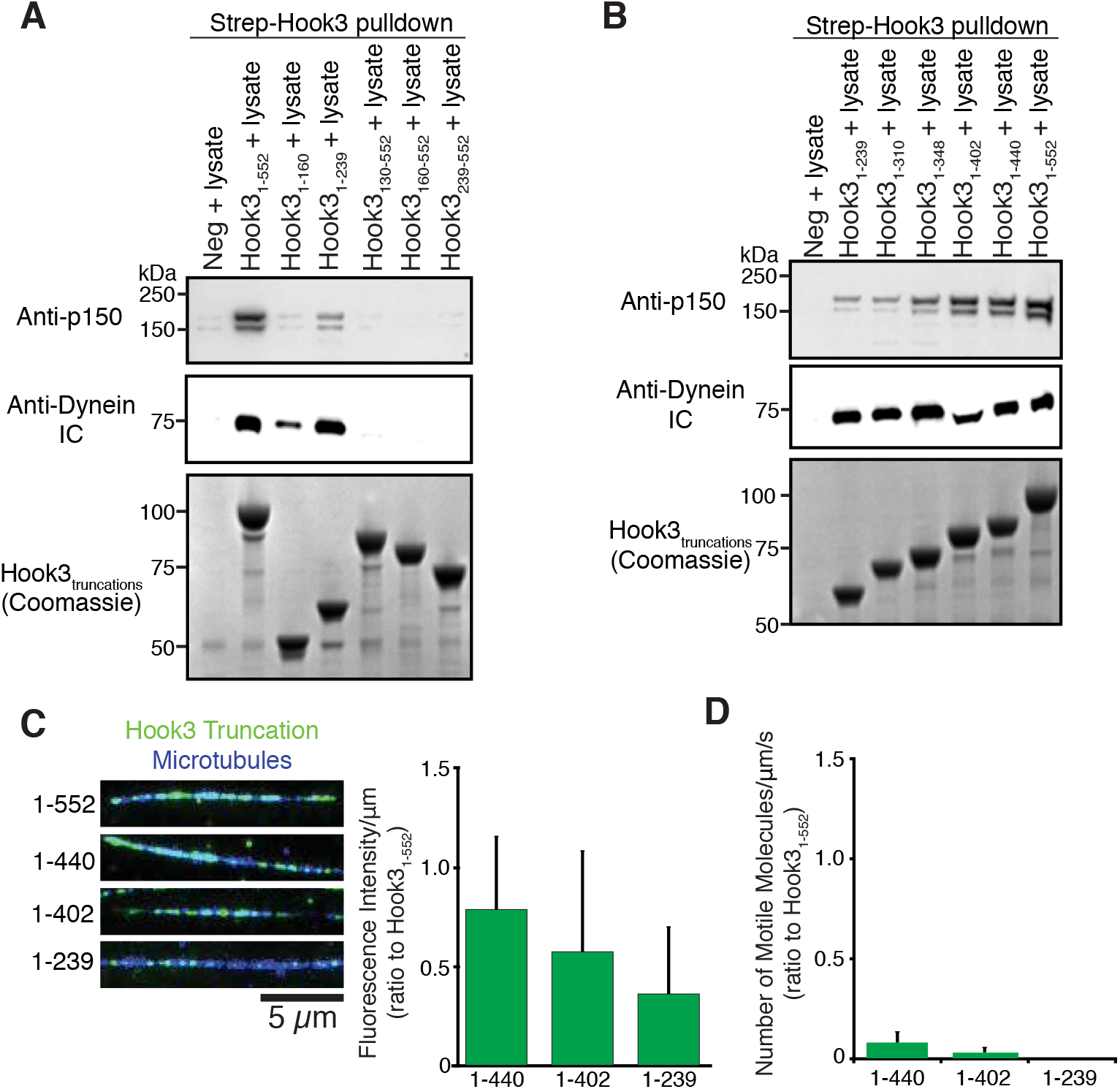
**Hook3 truncations that bind dynein-dynactin are not sufficient for motility A)** Truncations of strepII-Hook3 were tested for binding to endogenous dynein-dynactin in porcine brain lysate as in Fig. 4 B. Representative of three independent experiments. **B)** C-terminal strepII-Hook3 truncations were tested for binding porcine brain dynein-dynactin as in (A). The intermediate chain (IC) band in the lane for Hook3_1-402_ is skewed because the IC and this Hook truncation run at the same molecular weight. Representative of two independent experiments. **C)** The indicated truncations of SfGFP-Hook3 were incubated with affinity-purified human dynein-dynactin and fluorescence binding to surface-immobilized microtubules was assessed in the absence of ATP; overlay shows Hook3 in green and microtubules in blue (images are displayed using the same brightness and contrast). The fluorescence quantification for each condition is shown (mean fluorescence intensity (arbitrary units) of Hook3 per µm of microtubule). For each condition, >30 microtubules were quantified, and two replicate experiments were performed on different days (mean ± SD, with the SD representing the variation in the intensity/µm measured for all microtubules per construct). The scale bar represents 5 µm **D)** Hook3 constructs were tested for their ability to activate motility of the dynein-dynactin complex in the presence of ATP (see Fig. 4 C). The ratios of the shorter constructs to Hook3_1-552_ were calculated from side-by-side experiments performed on the same day. Shown are the mean ± SD of the ratios from two independent experiments performed on different days.

Since Hook3_1-239_ did not bind as much dynactin as Hook3_1-552_, we next investigated dynein and dynactin binding with a series of different length Hook3 constructs ending at residues 310, 348, 402, or 440 (residues chosen based upon the low probability of perturbing the structure of coiled coil 2, Lupas et al., 1991, Fig. S5 B). Lengthening the coiled coil domain from residue 239 to 552 did not significantly change the amount of dynein that is pulled down with Hook3 from the brain lysate (Fig. 5 B, Fig. S5 A). However, lengthening the coiled coil resulted in a progressive increase in the amount of interacting dynactin (Fig. 5 B, Fig. S5 A). These results suggest dynactin does not help to stabilize the Hook3-dynein LIC1 interaction. However, a longer Hook3 coiled coil is able to increase the affinity of dynein-Hook3 for dynactin.

We next investigated the microtubule binding ability and motility of the dynein-dynactin complex with Hook3_1-239_, Hook3_1-402_, Hook3_1-440_ and Hook3_1-552_. In this experiment, dynein and dynactin were first purified by affinity chromatography (see Methods) and then incubated with these truncated Hook proteins. In the absence of ATP, all truncated Hook3 proteins bound to microtubules by forming complexes with dynein or dynein-dynactin complexes with the longer constructs being somewhat more effective (Fig. 5 C). The Hook3 truncations did not bind microtubules in the absence of dynein-dynactin (Fig. S5 C). Surprisingly, in the presence of ATP and dynein-dynactin, all of the truncations induced poor single molecule motility compared to Hook3_1-552_ (Fig. 5 D). Hook3_1-239_ produced no processive motility at all, and even the longer constructs Hook3_1-402_ and Hook3_1-440_ produced very few motile events (Fig. 5 D, Fig. S5 D). The few complexes that were motile with Hook3_1-402_ and Hook3_1-440_ exhibited similar velocities to Hook3_1-552_ (Fig. S5 E). Therefore, truncations shorter than Hook3_1-552_, even though they can bind both dynein and dynactin, do not effectively produce complexes that can engage in processive motility. These findings indicate that the region of the coiled coil between aa 400-552 of Hook3 is required for robust activation of motility of the dynein-dynactin-Hook3 complex.

## Discussion

In this study, we have delineated the minimal binding regions for Hook3 that are required for three activities: 1) binding to the LIC1 C-terminal domain (Hook3_1-160_), 2) bringing dynein and dynactin together in a complex (Hook3_1-239_), and 3) producing a dynein-dynactin complex that engages in robust processive motility (Hook3_1-552_). Together, these results suggest a model for how cargo adaptors might regulate the minus-end-directed motility of dynein-dynactin.

Our work provides structural insights into how Hook3 binds to dynein. We previously found that the C-terminal half of LIC1 is the docking site for several cargo adaptors (Schroeder et al., 2014), and we show here that Hook3 binds to this same region of the LIC. Helix H, which extends from the CH domain, plays a key role in the LIC interaction, and our structure-function studies reveal two patches of residues in helix H (I136/I139/M142 and Q147/I154) that are involved in the interaction. These residues are highly conserved among all Hook isoforms, and thus it is likely that all Hook gene products bind LIC with a similar mechanism. Interestingly, in the unpublished NMR structure of the mouse Hook1 domain (PDB 1WIX), helix H is bent and the residues described above are sequestered in the middle of this bent conformation of helix H. Thus, based upon these two Hook domain structures, we speculate that helix H may be capable of undergoing a conformational change that could regulate its interaction with the dynein LIC. We speculate that Hook3_1-239_ may be able to bind LIC1 better than Hook3_1-160_ because this longer construct might shift a conformational equilibrium of helix H towards its extended form.

While the minimal Hook domain aa 1-160 binds the dynein LIC, it does not appear to be sufficient to recruit the dynactin complex. The first coiled coil of Hook3 (aa 160-239) enables dynactin binding and the additional coiled coil sequence further enhances this interaction. A cryo-EM study revealed that the 270-residue coiled coil of another cargo adaptor (BicD2) interacts along the groove of the Arp1 filament of dynactin and also mediates an interaction with the dynein heavy chain (Urnavicius et al., 2015). Similar to BicD2, Hook3’s coiled coils may sit in the groove of the Arp1 dynactin filament and promote an interaction between dynein and dynactin. Hook3_1-440_, for example, may have ~270 residues of coiled coil. However, the coiled coil of Hook3 (aa 160-552) alone is insufficient for stabilizing the tripartite complex, indicating that the Hook3-LIC1 interaction is also required. Supporting this conclusion, single point mutations in Hook3 (either Q147A or I154A) that abrogate LIC1 binding also completely block the ability of Hook3 to form a dynein-dynactin complex. Thus, multiple protein-protein interfaces of the adaptor Hook3 with the dynein heavy chain, LIC1, and dynactin appear to be required to form a stable tripartite motor complex.

We also show that the C-terminal region of our Hook3 construct is required for robust activation of dynein motility. A number of possible models could explain how this additional coiled coil-containing region converts an inactive dynein-dynactin-adaptor complex (e.g. one formed by Hook3_1-239_) into an active processive motor (one formed by Hook3_1-552_). First, a certain length of Hook3 bound along the dynactin Arp1 filament may be required to induce an allosteric conformational change in the dynein heavy chains to release them from an inhibited state (Chowdhury et al., 2015; Urnavicius et al., 2015). For example, an inhibitory state of dynein may exist in which the two motor domains are stacked, necessitating the separation and alignment in the same direction to become active (Torisawa et al., 2014). An alternative and not mutually exclusive model involves the allosteric regulation by Hook3 of the N-terminal CAP-Gly domain of dynactin’s p150 subunit. The p150 subunit regulates dynein motility (Kardon et al., 2009; McKenney et al., 2014; Tripathy et al., 2014), and p150’s CAP-Gly domain binds to the C-terminus of tubulin, an interaction that greatly enhances an initial microtubule binding encounter of dynein-dynactin-BicD2 that leads to processive movements (McKenney et al., 2016). However, dynactin alone exhibits minimal binding to microtubules, suggesting that it is in an autoinhibited state (Kardon et al., 2009; McKenney et al., 2014). This finding agrees with a dynactin cryo-EM structure showing that the junction between CC1A and CC1B in p150 is positioned near the pointed end of the Arp1 filament; in this folded conformation the CAP-Gly and CC1A domains are unlikely to be accessible to the microtubule (Urnavicius et al., 2015). In a lower resolution structure of the dynein-dynactin-BicD2 complex, the C-terminus of a 270 aa coiled coil of BicD2 is located at the pointed end of the dynactin Arp1 filament (Urnavicius et al., 2015). We speculate that the C-terminal end of our motility-inducing Hook3 construct (aa 400-552) may reach to and perhaps beyond the pointed end of the Arp1 filament and somehow act to dislodge CC1A-CC1B from the backbone of the Arp1 filament. The release of p150 may enable this subunit to extend fully into an active conformation, enabling access to the microtubule (Fig. 6).

**Figure 6.**
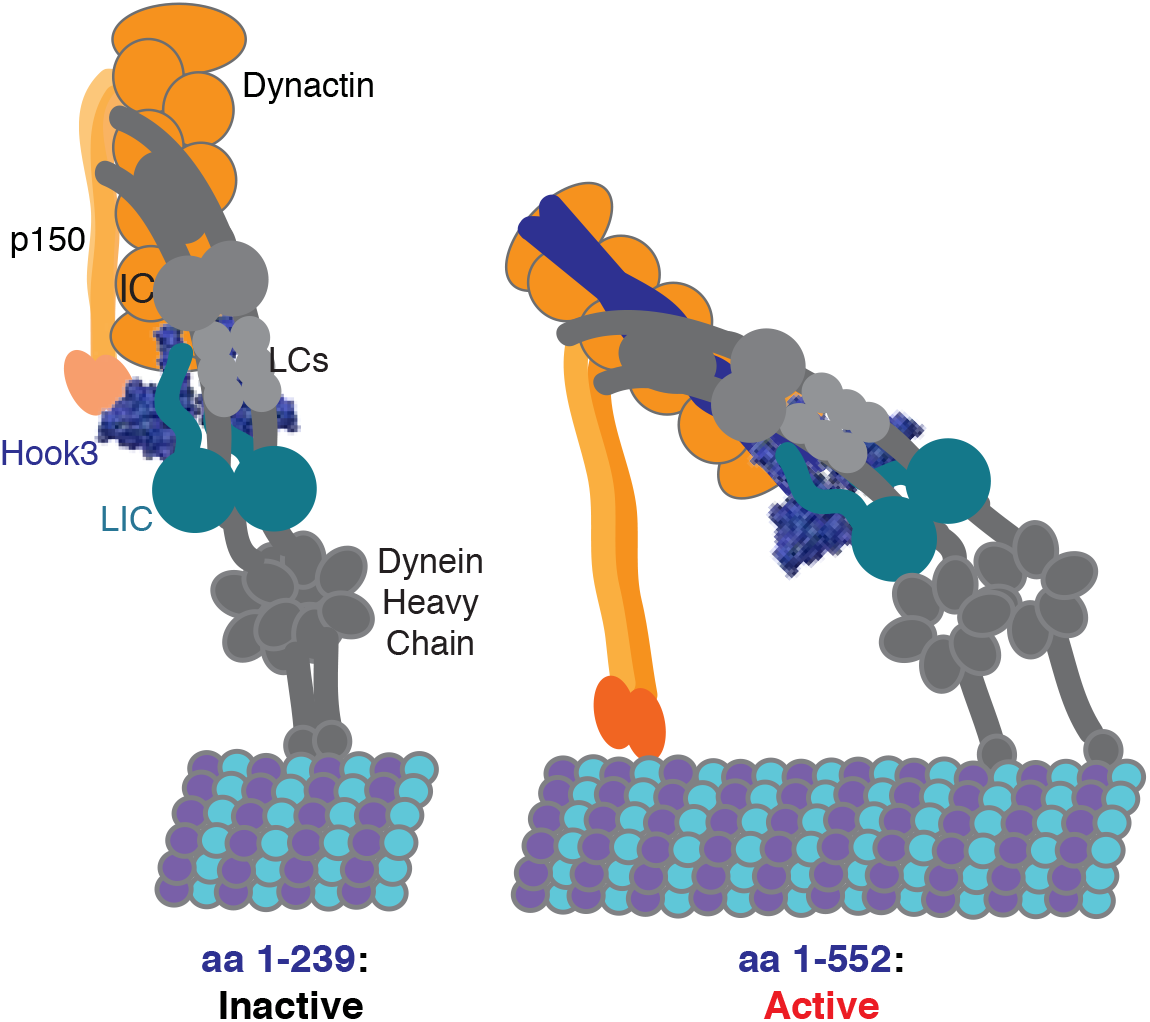
**Model of assembly and activation of the dynein-dynactin-Hook3 complex** Short Hook constructs (e.g. Hook3_1-239_) are able to assemble the tripartite motor complex by binding the LIC1 C-terminal domain and part of the dynactin Arp1 filament and dynein heavy chain. However, the complex is inactive for motility. We speculate that the longer coiled coil of Hook3_1-552_ releases the CAP-Gly domain of p150 from an autoinhibited state to enable its binding to microtubules, thus enhancing the initiation of processive motility. A change in the orientation or other allosteric change in the motor domains, based upon work of Urnavicius et al. (2015) and Torisawa et al. (2014), also might be promoted by the longer Hook constructs. The illustration of the dynein-dynactin complex is based upon work by Urnavicius et al. (2015) and Chowdhury et al. (2015).

While our data reveal an important role of the LIC in Hook3-mediated dynein motility, a number of questions remain unanswered. First, it is unknown whether the LIC acts as a passive tether for linking the motor domain to cargo adaptors or whether it also undergoes a conformational change that plays an active role in eliciting dynein motility. Second, it remains to be determined whether other cargo adaptors that interact with LIC1 (e.g. FIP3, RILP, BicD2) do so through similar or different mechanisms to Hook proteins. Among these adaptors, the CH domain and helix H of Hook proteins appear to be unique. Third, we also do not know how many cargo adaptors use the LIC for dynein activation and which ones might work through LIC-independent mechanisms. Differences in activation among the adaptors might allow for further regulation of dynein-based cargo transport. Many of these important questions can be addressed through structural and functional studies of multiple types of cargo adaptor proteins interacting with dynein and dynactin.

## Materials and Methods

### Molecular Biology

The cDNA of Hook3 was obtained from a human cDNA library made from mitotic RPE-1 cells, and all Hook3 constructs were cloned into vector pET28a with an N-terminal 6x His-strepII-superfolder GFP (sfGFP) tag. All human LIC1 (NM_016141.3) constructs were cloned into pGEX6P1, which included an N-terminal GST tag and a C-terminal strepII tag. Truncations were based on both secondary structure prediction (Drozdetskiy et al., 2015) and coiled coils prediction (Lupas et al., 1991). To dimerize the Hook3_1-160_ construct, the GCN4 sequence (Harbury et al., 1993) was added to the C-terminus. The 29 amino acid sequence was VKQLEDKVEELLSKNAHLENEVARLKKLV. Full-length human FIP3 (accession AB383948) was cloned into pET28a with a strepII-SNAP tag and was used for the purification of the dynein-dynactin complex from porcine brain lysate.

### Protein Purification

All human Hook3 constructs were transformed into the *E. coli* strain BL21 RIPL and expression was induced with 0.5 mM IPTG at 37ºC for 3-6 hr. Bacterial pellets were resuspended with lysis buffer (25 mM Hepes pH 7.8, 150 mM NaCl, 2 mM TCEP, 1 mM PMSF, and a protease inhibitor cocktail—Roche, 1 tablet per 50 ml) and lysed using an Emulsiflex press (Avestin). The lysate was clarified by centrifugation at 40,000 × g for 30 min and Hook3 was purified using Strep-Tactin Superflow Plus resin (QIAGEN). The agarose was then washed with lysis buffer (excluding the Roche protease inhibitor cocktail) at approximately 20X the resin volume, and then the purified protein was eluted with 3 mM desthiobiotin. The protein was concentrated and flash frozen. Thawed protein was then further purified by gel filtration with a Superose 6 10/300 GL column (GE healthcare) or a Superdex 200 10/300 GL (GE healthcare). The gel filtration buffer was 30 mM Hepes pH 7.8, 150 mM NaCl, 2 mM MgCl_2_, 5% glycerol and 2 mM TCEP. The Hook3-containing fractions were pooled, concentrated and flash frozen. StrepII-SNAP-FIP3 was purified the same way as Hook3 except the lysis buffer included 25 mM Tris pH 8.5.

GST-LIC1-strepII constructs (full-length and truncations) were expressed as performed with the Hook3 constructs followed by lysis with 50 mM Tris pH 7.4, 150 mM NaCl, 5 mM TCEP, 1 mM PMSF, and a protease inhibitor cocktail—Roche, 1 tablet per 50 ml). The protein was then purified using either glutathione agarose 4B (USB) or Strep-Tactin Superflow Plus resin (QIAGEN). After extensive washing and elution with either 10 mM reduced glutathione at pH 7.4 (for glutathione agarose) or 3 mM desthiobiotin (for Strep-Tactin resin), the protein was gel filtered using a HiPrep 16/60 Superdex S-200 HR column (GE Healthcare) in 20 mM Tris pH 7.4, 50 mM NaCl and 2 mM TCEP.

### Pulldowns

Clarified porcine brain lysate was used to test the binding of endogenous dynein-dynactin to Hook3 constructs and was prepared as previously done (McKenney et al., 2014). For each dynein-dynactin pulldown, 500 µl of porcine brain lysate in buffer A (30 mM Hepes, pH 7.4, 50 mM K-acetate, 2 mM Mg-acetate, 1 mM EGTA, 10% glycerol) was combined with 60 µl of 50% Strep-Tactin sepharose slurry (GE Healthcare), 0.1% NP40, 5 mM DTT, and 1 mM PMSF. SfGFP-tagged Hook3 constructs were added at 200-400 nM to the brain lysate and resin and incubated for 1-2 hr at 4ºC. The resin was pelleted and washed 5X in 500 µl buffer A including 0.1% NP40 and 5 mM DTT. After the final wash, the resin was resuspended in 50 µl of loading buffer and an equal volume of the samples was resolved on NuPAGE gels (Invitrogen). All dynein-dynactin pulldowns were repeated two or three times on separate days starting from frozen brain lysate.

To test the binding of human LIC1 to Hook3 constructs, 200 nM GST-LIC1-strepII (full-length or truncations), was incubated with 20 µl glutathione resin in a 300 µl volume of buffer (50 mM Tris, pH 7.4, 100 mM NaCl, 5 mM TCEP, 0.1% Tween, and 2 mg/ml BSA). After washing the resin extensively, 300 µl of 200 nM sfGFP-Hook3 construct was added and incubated for 1 hr at 4ºC. The resin was washed extensively and then resuspended in 20 µl of 1X loading buffer. Samples were resolved on NuPAGE gels. All pulldowns with purified LIC1 and Hook3 proteins were repeated two or three times on separate days.

### Western blot analysis

After samples were resolved by SDS-PAGE, the samples were transferred to nitrocellulose membranes with the iBlot Gel Transfer Device (Invitrogen). Membranes were blocked with 5% milk in TBS and 0.1% Tween (TBST) and probed at room temperature with primary antibody, which included rabbit anti-GFP (Abcam, 1:1000), mouse anti-GST (Thermo Fisher Scientific, 1:1000), mouse anti-dynein intermediate chain (clone 74.1, Millipore, 1:1000), and mouse anti-p150 (BD Biosciences, 1:250). Membranes were then washed three times with TBST and incubated with anti-mouse-800 or anti-rabbit-680 (Molecular Probes, 1:10,000) for 45 min −1 hr at room temperature. Blots were visualized by an Odyssey Clx Infrared Imaging System (LI-COR).

### Crystallization and Structure Determination

The LIC1 C-terminal half (LIC_389-523_) and GST-Hook3_1-160_ were purified with glutathione agarose resin 4B (USB) and cleaved from the resin using purified GST-tagged human rhinovirus 3C protease and overnight incubation at 4ºC. After the GST tag was cleaved, the two proteins were combined at an equimolar ratio and incubated on ice for 30 min. The proteins were gel filtered using a HiPrep 16/60 Superdex S-200 HR column (GE Healthcare Life Sciences) into the following buffer: 10 mM Tris pH 7.4, 25 mM NaCl, 2 mM TCEP. Fractions containing both proteins were concentrated to approximately 20 mg/ml and hanging drop vapor diffusion experiments were setup using 96-well crystal screens (QIAGEN) at room temperature. Native crystals grew from a reservoir solution containing 2 M sodium formate, 0.1 M sodium acetate pH 4.6 (JCSG screen Core III, QIAGEN). The crystals were cryoprotected with the addition of 35% glycerol to the crystallizing well solution and were flash cooled by plunging in liquid nitrogen.

Native diffraction data were collected at beamline 8.3.1 at the Advanced Light Source (Lawrence Berkeley National Laboratory), and the dataset was indexed and integrated in P 2 21 21 using XDS (Kabsch, 2010). The structure was solved by molecular replacement using an ensemble of 20 superimposed NMR models from PDB structure 1WIX using Phaser (McCoy et al., 2007). The Phaser scores for the best solution were modest (RFZ=4.8 TFZ=6.4) and the initial electron density maps were noisy and discontinuous. Density modification and chain tracing with SHELXE (Sheldrick, 2010) resulted in an easily interpretable map and a poly-Alanine model that was further improved using phenix.autobuild (Terwilliger et al., 2008). Multiple rounds of model building and refinement were done using Coot (Emsley and Cowtan, 2004) and phenix.refine (Adams et al., 2010). The data collection and refinement states are presented in Table I, and the PDB accession code is 5J8E.

### Purification of dynein-dynactin from human RPE-1 cells

RPE-1 cell lysate was prepared as previously described (McKenney et al., 2014). The lysate was centrifuged at 266,000 × g for 10 min at 4ºC and a final concentration of 5 mM DTT, 0.1% NP40, and 1 mM PMSF was added before use. The lysate was incubated with purified strepII-SNAP-FIP3 on Strep-Tactin sepharose (GE Healthcare). After incubation at 4ºC for 1-2 hr, the resin was thoroughly washed with buffer A (30 mM Hepes, pH 7.4, 50 mM K-acetate, 2 mM Mg-acetate, 1 mM EGTA, 10% glycerol, 5 mM DTT, 0.1% NP40 and 1 mM PMSF) and then resuspended in buffer A with 300 mM NaCl to release dynein-dynactin from resin-bound FIP3. After incubating on ice for 10 min, the high-salt slurry was centrifuged through a 0.2 µm filter to remove the resin. Then an equal volume of 50% Strep-Tactin sepharose slurry was added to the elution to bind any strepII-FIP3 that may have released from the resin during the high-salt incubation. After incubating on ice for 10 min, the slurry was once again filtered, and the final solution was diluted with buffer A for a final concentration of 200 mM NaCl. Sucrose was also added at a final 6% concentration, and the affinity purified dynein-dynactin was flash frozen for single molecule imaging.

### Single molecule imaging

#### Preparation of Microtubules

Tubulin was purified from porcine brain and labeled (fluorescently or with biotin) as previously described (Castoldi and Popov, 2003). To polymerize microtubules, unlabeled tubulin was combined with biotin-labeled tubulin and fluorescent tubulin (640 nm fluorescence) at a ratio of approximately 10:2:1, respectively, in BRB80 (80 mM PIPES, 1 mM EGTA, 1mM MgCl2) and 5 mM GTP. After incubating for 10 min at 37ºC, taxol was added at a final concentration of 20 µM. To remove unpolymerized tubulin, the microtubules were layered over a 25% sucrose cushion and centrifuged at 65,000 × g for 5 min at 22ºC.

#### Preparation of dynein-dynactin-Hook3 complexes

A 30 µl reaction consisting of 10 nM sfGFP-tagged Hook3 and 5 µl of ~0.15 mg/ml native dynein-dynactin purified from RPE-1 cells was incubated in 30 mM HEPES pH 7.4, 50 mM K-acetate, 2 mM Mg-acetate, 1 mM EGTA, 10% glycerol, 0.1 mg/ml BSA, 0.5% pluronic acid F-127, 0.2 mg/ml kappa-casein, and a trolox/PCA/PCD scavenging system (Dave et al., 2009).

#### Total Internal Reflection Fluorescence (TIRF) Microscopy

Flow chambers (volume ~10 µl) were constructed using double-sided stick tape and acid-washed coverslips as described (Tanenbaum et al., 2013). The chambers were prepared with immobilized fluorescent microtubules by coating the chamber in the following sequence of solutions: 10 µl 5 mg/ml BSA-biotin (Thermo Scientific), 20 µl BC buffer (BRB80, 1 mg/ml BSA, 1 mg/ml casein, 0.5% pluronic acid F-68, pH 6.8), 10 µl 0.5 mg/ml streptavidin (Vector labs), 20 µl BC buffer, and finally 10 µl of a 1:10 dilution of microtubules (as prepared above). Microtubules were washed with the assay buffer and then a 1:10 dilution of the dynein-dynactin-Hook3 complex described above was added to the flow chamber in the presence of 1 mM ATP.

Movies were acquired with a Nikon Eclipse TE200-E microscope equipped with an Andor iXon EM CCD camera, a 100X 1.49 NA objective, and Micromanager software (Edelstein et al., 2010). A 491 laser (at 75% laser power) and a 640 nm laser (at half max laser power) were used to image sfGFP-Hook3 (100 ms exposure) and fluorescently labeled microtubules (50 ms exposure), respectively. Several 6 min movies (1 or 2 s intervals of image acquisition) were acquired at room temperature per flow chamber per construct. Molecules that moved >1 µm were scored as processive. Velocities were quantified by making kymographs in ImageJ (NIH).

## Accession Code

The Protein Data Bank Accession Code is 5J8E.

## Summary of Supplemental Material

Fig. S1 shows the complete gels and relative inputs for Fig. 1’s pulldowns, and it shows the cogel filtration of LIC_1389-523_ and Hook3_1-239_. Fig. S2 shows the alignment of human Hook3 and mouse Hook1 (PDB 1WIX), and it displays the conservation of the hook domains of the human Hook isoforms. Fig. S3 shows the sequence alignment used to map the conservation of Hook domains onto the Hook3_1-160_ structure in Fig. 2 B. Fig. S4 presents the gel filtration chromatograms of all Hook3 proteins with triple mutations and representative kymographs for the Hook3 single point mutants, related to Fig. 4. Fig. S5 displays the predicted coiled coils of Hook3, Hook3’s interaction with microtubules in the presence and absence of dynein-dynactin, and representative kymographs for Fig. 5 C.

**Figure S1.**
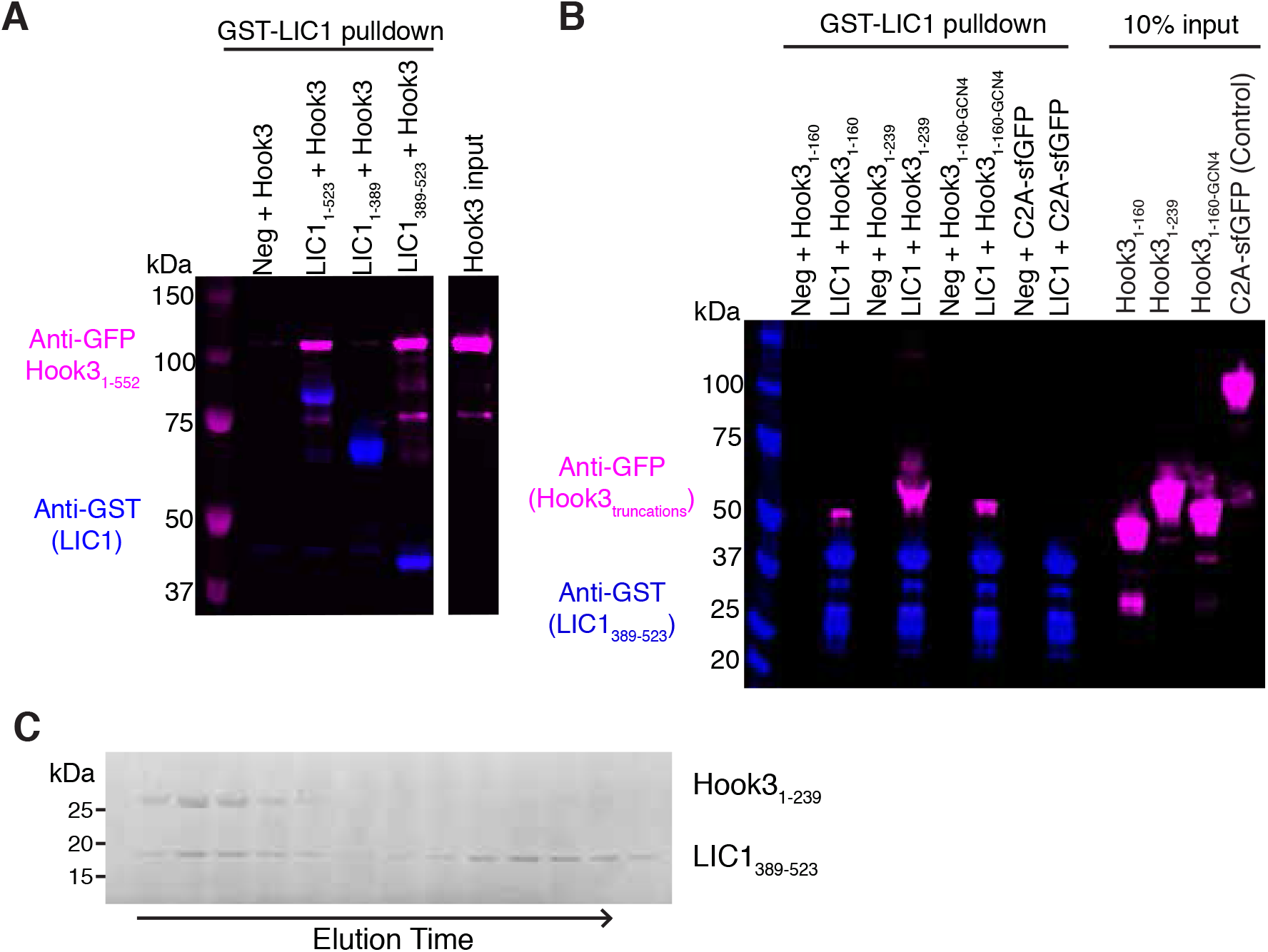
**Determination of the minimal binding regions of the LIC1-Hook3 complex A)** The gel for Fig. 1 B is displayed with the corresponding Hook3 input to indicate the amount of Hook3_1-552_ used in the assay. **B)** The full gel for Fig. 1 C is displayed with the corresponding Hook3 input to indicate the amount of Hook3_1-160_, Hook3_1-239_, and Hook3_1-160-GCN4_ used in the assay. An unrelated protein (sfGFP-tagged membrane binding domain C2A of synaptotagmin-I, Hui et al., 2011) was also tested for binding to the LIC as a negative control. **C)** Human LIC1_389-523_ was incubated with human Hook3_1-239_ on ice for 30 min with both proteins at 20 µM. The protein mixture was then subjected to gel filtration on a S200 10/300 GL column (GE Healthcare) with 25 mM Tris pH 7.4, 100 mM NaCl, and 2 mM TCEP. Fractions were analyzed by SDS-Page (Coomassie stain). A portion of the LIC1_389-523_ co-elutes with Hook3_1-239_ (left side of the gel).

**Figure S2.**
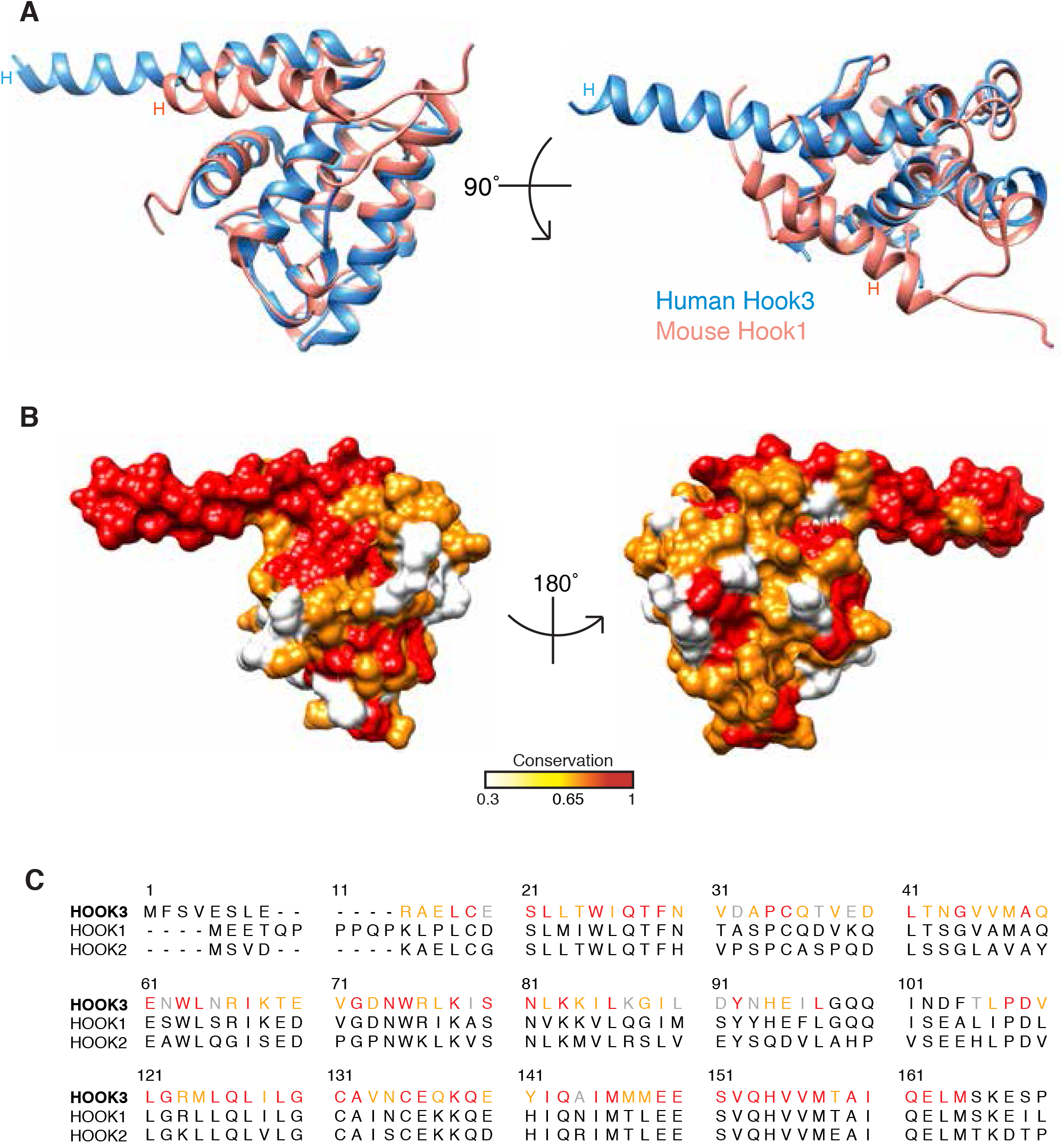
**Conservation of the Hook domain A)** The crystal structure human Hook3 (aa 9-158) is aligned with the solution structure of mouse Hook1 aa 1-164 (PDB 1WIX) using Chimera. Two views are presented to show the bent (mouse solution structure) and extended (human crystal structure) structures of helix H. **B)** The Hook domain sequences of human Hook1, 2 and 3 were aligned using MAFFT version 7 (Katoh and Standley, 2013) and the conservation of residues is mapped onto the crystal structure of the human Hook3 domain. The surface conservation is displayed and the percent of conservation is depicted as a gradation of red with red being 100% conserved. **C)** The sequence alignment used in (B) is presented with a similar color scheme in the Hook3 sequence to show conservation. Red depicts 100% conserved, whereas the least conserved residues (30%) are shown as gray.

**Figure S3.**
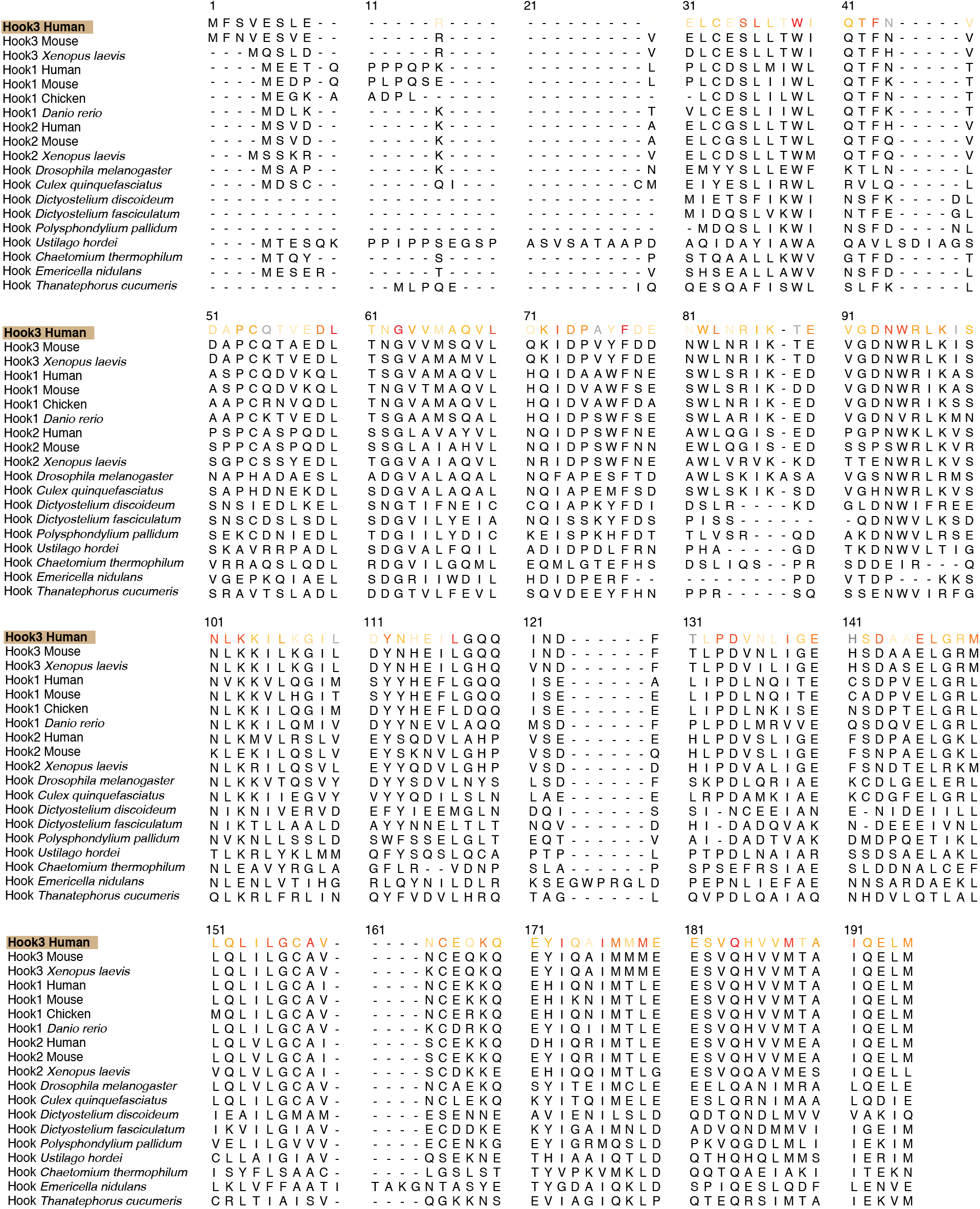
**Sequence alignment of the Hook domain from diverse species** Nineteen Hook domains (~160 residues) ranging from fungi to metazoans were aligned using MAFFT version 7 (Katoh and Standley, 2013) and viewed in Chimera. Conservation is depicted in the human Hook3 sequence with a gradation of red similar to Fig. 2 B. Dark red is 100% identical. The least conserved residues are shown in gray.

**Figure S4.**
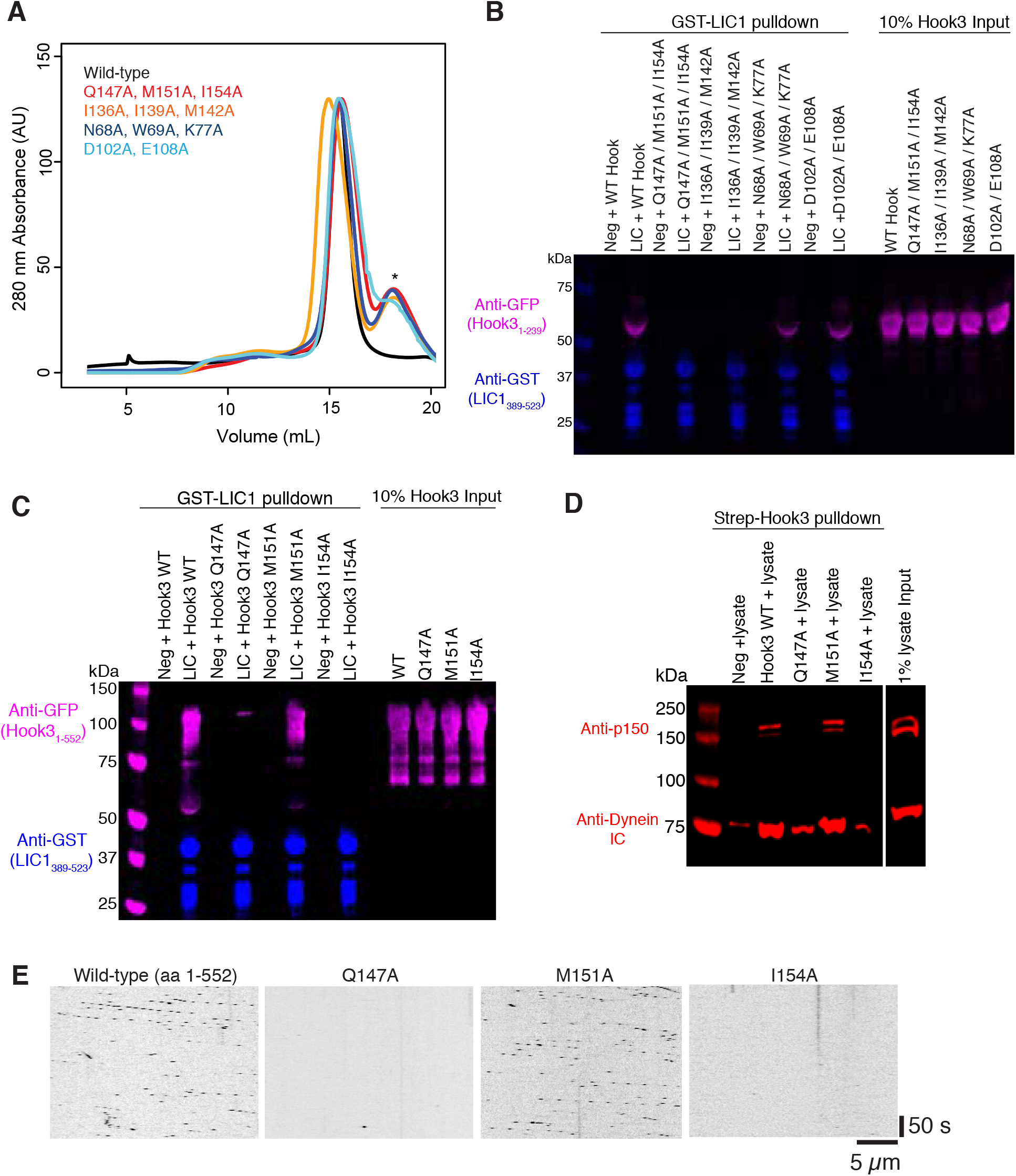
**Hook3’s conserved residues are critical for the assembly of the tripartite dynein complex A)** Wild-type Hook3 and the triple and double mutants were each gel filtered on a Superose 6 10/300 GL column with 30 mM HEPES pH 7.4, 150 mM NaCl, 2 mM MgCl2, 5% glycerol, and 2 mM TCEP. The chromatograms are displayed with wild-type in black. The asterisk denotes sfGFP alone as a protease cleavage product. Wild-type did not have sfGFP contamination because it had previously been gel filtered. The fractions at ~16 ml were collected and concentrated for experiments. **B)** The full gel for Fig. 3 B is displayed with the corresponding Hook3 input to indicate the amount of Hook3 mutants used in the assay. **C)** The full gel for a repeat of the experiment shown in Fig. 4 A is displayed. In this experiment, the Hook3 input was double the concentration used in Fig. 4A’s pulldown to show the Hook3 mutant Q147A binds LIC1 slightly. **D)** The full gel for Fig. 4 B is displayed with the corresponding input of endogenous dynein and dynactin present in the porcine brain lysate. **E)** Wild-type sfGFP-Hook3 and the single-point mutants Q147A, M151A, and I154A (all aa 1-552 of Hook3) were tested for activation of dynein-dynactin motility. Kymographs were made to assess motility, and representative kymographs are displayed using the same brightness and contrast. The interrupted lines are due to the acquisition parameters (100 ms exposure every 2 s to minimize photobleaching). The scale bars represent 50 s and 5 µm. The quantification of all kymographs is displayed in Fig. 4 C.

**Figure S5.**
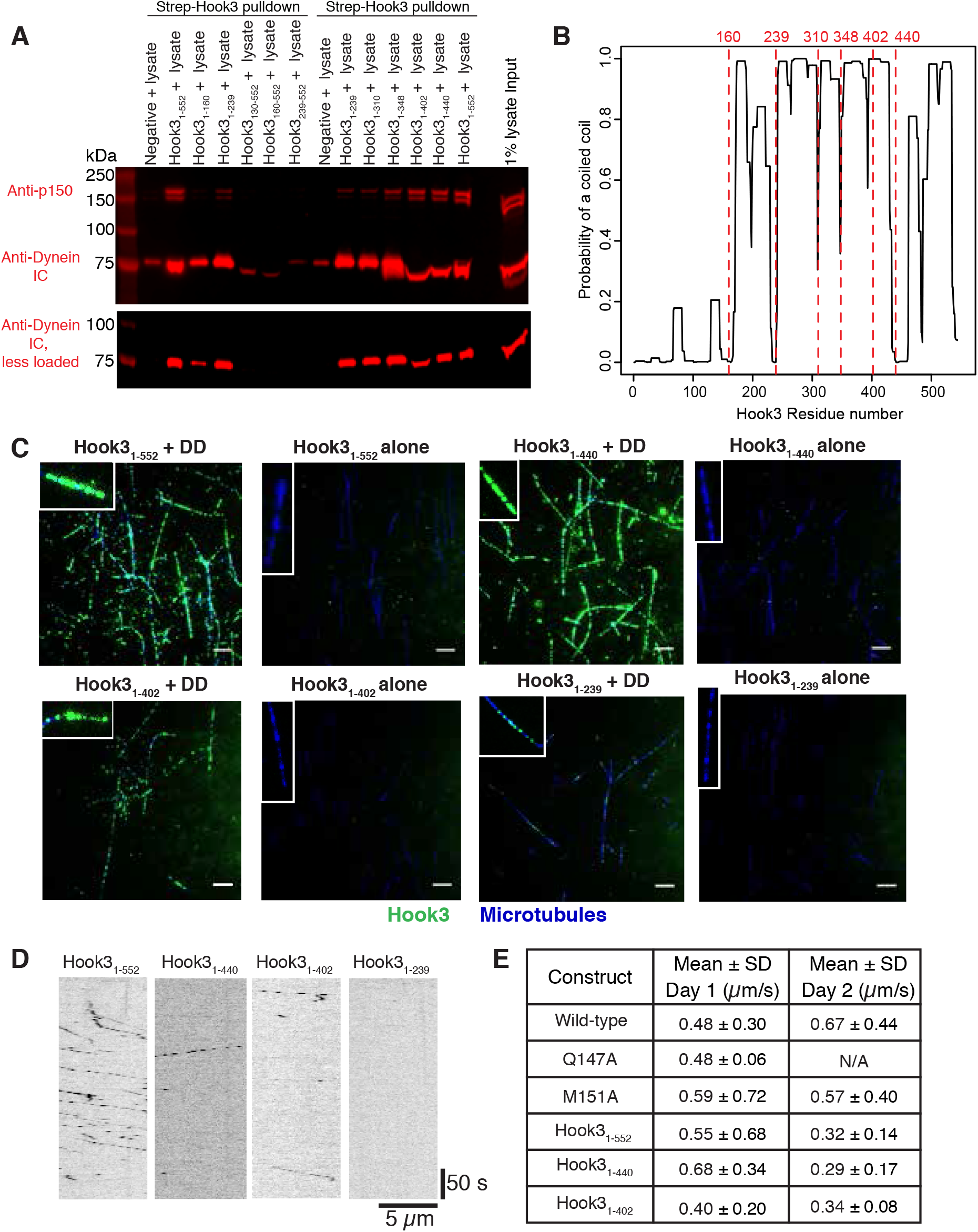
**Some Hook3 truncations can bind dynein-dynactin but have limited motility A)** The full gel for Fig. 5 A and B, which were done simultaneously in this experiment, is displayed along with the corresponding input of endogenous dynein-dynactin present in the porcine brain lysate. The amount of the samples loaded for dynactin detection was 25 µl, which led to distortion of the bands corresponding to lower molecular weights. To show the dynein intermediate chain undistorted, the samples were re-run with less volume (10 µl), shown below the first immunoblot. **B)** The amino acid sequence of Hook3_1-552_ was analyzed for probability of forming a coiled coil with the COILS server (Lupas et al., 1991) using a narrow window. The graph indicates the probability of a coiled coil versus the residue number of Hook3_1-552_. The truncations made at aa 160, 239, 310, 348, 402, and 440 are denoted with a red line and label. **C)** SfGFP-Hook3 truncations were incubated with or without human dynein-dynactin (DD) for 30 min on ice and then added to glass-immobilized microtubules. After incubating for 3 min, unbound complexes were washed away, and the microtubules were then viewed by TIRF microscopy to assess bound sfGFP-Hook3. sfGFP-Hook3 is displayed as green, and microtubules are blue. Brightness and contrast are the same between images showing a Hook3 truncation with and without dynein-dynactin. Insets show closer views of representative microtubules, and scale bars represent 5 µm. **D)** The truncations in Fig. 5 D were tested for activation of dynein-dynactin motility, and representative kymographs are shown for each truncation. The kymographs are displayed using the same brightness and contrast, and the scale bars represent 50 s and 5 µm. **E)** The mean velocities ± SD are displayed for each construct for two independent experiments conducted on different days. The number of molecules measured for wild-type Hook3_1-552_ and single point mutants on day 1 and day 2 are as follows: wild-type n = 146 and n = 135; Q147A n = 2 and n = 0; M151A n = 199 and n = 203. I154A had no motile molecules. The number of molecules measured for Hook3 truncations on day 1 and day 2 are as follows: Hook3_1-552_ n = 199 and n = 401; Hook3_1-440_ n = 28 and n = 16; Hook3_1-402_ n = 12 and n = 3. Hook3_1-239_ had no motile molecules.

## Acknowledgments

We thank Damian Ekiert for his help with crystallographic data acquisition and molecular replacement. We thank Walter Huynh for his help in optimizing the single molecule imaging and Richard McKenney for his advice on purifying dynein-dynactin. We also thank Nico Stuurman for his help with the microscopy and the rest of the Vale lab for useful discussions.

## Abbreviations List

CC: coiled coil
LIC: light intermediate chain
Neg: negative
sfGFP: superfolder-green fluorescent protein
WT: wild-type

